# Changes in the transcriptome and long non-coding RNAs but not the methylome occur in human cells exposed to *Borrelia burgdorferi*

**DOI:** 10.1101/2022.07.11.499513

**Authors:** Anne Berthold, Vett Lloyd

**Author notes:** Correspondence: Anne Berthold.

## Abstract

Lyme disease, caused by infection with members of the Lyme borreliosis group of *Borrelia* spirochete bacteria, is increasing in frequency and distribution worldwide. This spread is driven by the expansion of ticks, vectors of these pathogens. Epigenetics may be involved in the interaction between mammalian host, tick, and bacterial pathogen, but is still poorly understood. Next-generation sequencing (NGS) allows the study of host response at the transcriptomic and methylomic scale.

We tested the effect of the *Borrelia burgdorferi* strain B31 on a human primary cell line (HUVEC) and an immortalized cell line (HEK-293) for 72 h, a time consistent with the duration of tick feeding and host cell exposure to *B. burgdorferi*. RNA and DNA were extracted from cells and used for RNA-seq and Enzymatic-Methyl-seq (EM-seq). Differential expression and Reactome pathway enrichment analysis were performed. More significantly differentially expressed genes (a total of 69) were found in HUVECs compared to HEK-293 (a total of 8). *Borrelia burgdorferi* exposure significantly induced genes in the interferon, cytokine, and other immune response signaling in HUVECs. In HEK-293, pre-NOTCH processing in Golgi was identified as a significant pathway. Other significant genes suggest extracellular matrix binding and interaction in HEK-293. Data comparison with other transcriptomic studies of human cells exposed to *B. burgdorferi* revealed a small overlap in genes identified in HUVEC, but no overlap for HEK-293. No significant methylation changes were detected in HUVECs or HEK-293 exposed to *B. burgdorferi*. However, two long non-coding RNAs and a pseudogene were deregulated in response to *B. burgdorferi* in HUVEC suggesting that other epigenetic mechanisms may be initiated by infection.

This is the first study examining the transcriptome and methylome of human cells exposed to *B. burgdorferi* for 72 h and so contributes to the description of early-stage *B. burgdorferi* infection at the cellular level.

## 1. Introduction

Zoonotic diseases are the focus of a global research effort, in part due to the recent pandemic. Ticks are notorious vectors for transferring zoonotic pathogens to humans and other animals. One tick-borne disease is Lyme disease, which is caused by the spirochete *Borrelia burgdorferi* and related species [1]. As the disease is spreading across temperate regions of the globe including North America and Europe, it is becoming an emerging public health concern [2]. Climate change is one of many factors contributing to tick population expansion and success [3–5], thus encounters between humans and ticks, and tick-borne pathogens, are expected to increase in the near future. The incidence of Lyme disease has increased over the past decades, as evidenced by surveillance data from several European countries and the United States [6–14], and the World Health Organization (WHO) office in Europe [15]. The annual Lyme disease case numbers are estimated to be ∼65,000 to 85,000 cases in Europe [15, 16] and ∼300,000 cases in the USA [9, 10]. Even so, tick-borne diseases are likely under-reported as access to testing varies within and between countries, the sensitivity and specificity of the most common serological tests are imperfect and geographically variable, and case definitions varied among countries [17–19].

*Borrelia* infection can cause Lyme disease, which is divided into three stages: early localized, early disseminated, and late disseminated infection [20]. Both local and disseminated multisystemic symptoms place a burden on patients, which is exacerbated in cases where there is a lack of risk recognition, accurate diagnosis, and treatment of the disease [21]. Transmission of tick-borne pathogens occurs primarily by injection as the tick feeds and tick salivary proteins modulate host defense mechanisms [22]. Studies with *B. burgdorferi* have shown that, depending on the cell or animal model, different signaling molecules in the immune response cascade are downregulated promoting pathogen survival and spread of the pathogen in the host [20, 23]. The symptomatology of the disease is manifold and varies from patient to patient [24]. Even after treatment, ongoing symptoms due to failure to clear the infection, cellular damage/alteration, or altered immune response can occur [25].

Epigenetic processes are central to the regulation of biological processes of an organism and their misregulation contributes to the development of diseases; immune function is no exception [26]. An organism may be exposed to various environmental factors in the short or long term, including bacterial pathogens [27]. These environmental factors leave epigenetic marks that complement the genetic code of the host to determine the level of gene expression and function [28]. Epigenetic mechanisms are complex, highly dynamic, and reversible yet can be transmitted to the next generation [29]. Major epigenetic mechanisms include chemical modifications of histones or DNA bases, the insertion of histone variants and remodeling complexes, and the action of non-coding RNAs (ncRNA) [28].

Embryogenesis and cell differentiation processes are a few examples where properly-timed epigenetic events, such as DNA methylation and histone modifications, often orchestrated by ncRNAs, take place and contribute to cell identity, differentiation, and the normal epigenetic landscape of cells and organisms [30]. A cell-type-specific spatial and temporal resolution of genome functions, including all epigenetic mechanism and their interplay, is essential for a detailed understanding of normal and pathophysiological states. Bacteria can alter the epigenome of the host they infect, reprogramming host genes and leading to alteration of host physiology, morphology, and behavior [31]. These adaptations have been selected to be beneficial to bacteria, allowing the bacteria to infect, persist, and spread within the host. For example, the bacterial pathogens *Anaplasma* manipulate physiological processes in their tick vectors by altering histone-modifying enzymes [32]. *Borrelia burgdorferi* methyltransferases have been characterized and regulate gene expression in these bacteria [33]. In astrocytes infected with *B. burgdorferi*, transcriptomic and epigenomic analyses showed that access to chromatin decreased during the course of infection, which was associated with anatomical and morphological changes and eventually stress mechanisms in the cells [34]. Thus, epigenetic mechanisms are likely to play an important role in *Borrelia* infection during host-pathogen interactions [31, 35].

Interactions between *Borrelia* and numerous mammalian cell types have been studied *in vitro* and *in vivo.* Studied cells include keratinocytes [36], fibroblasts [37–40], neuronal cells [41–45], and blood-derived cells [46–49]. This study focused on interactions between *B. burgdorferi* and human cells *in vitro*. We hypothesize that exposure of human cells to bacteria would lead to a change in transcription that might be due to altered DNA methylation. We used two cell models, one endothelial and one epithelial, as these cell types would be expected to respond differently. The epidermis, dermis, and the various cell populations therein, represent the first physical barriers that ticks and pathogens must overcome to establish infection [50]. Human umbilical vein endothelial cells (HUVECs) have previously served as primary cell models in several pathogen-host interaction studies [51, 52], to explore the localization of *Borrelia* in the cell [37,44,53], and to study transcriptional changes [54]. Human embryonic kidney cells (HEK-293) have been used to study the attachment of *B. burgdorferi* to extracellular matrix components such as negatively-charged polysaccharide glycosaminoglycans (GAG) and proteoglycans [41,43,55]. The blood meal of hard-bodied ticks *(Ixodidae*) which typically vectors *Borrelia* species, usually lasts for several days [56]. We therefore chose an exposure time of 72 h, more prolonged exposure than most other studies which have been restricted to exposure times between 2 and 24 h [36–38,40,42,44,45,47,49]. In addition to this prolonged exposure time reflective of *Borrelia*-host exposure during the blood meal, this time period might also encompass the internalization of *Borrelia* in immune and other cells in the early stages of the disease. We used live bacterial cells because it has been shown that in peripheral blood mononuclear cells live *B. burgdorferi* 297 induce a stronger inflammatory signal than bacterial lysate [49]. By studying both transcriptional and methylome changes in two different human cell types exposed to *B. burgdorferi* for 72 h, we expand knowledge of the cellular interface between the host and the tick-borne pathogen.

## 2. Results

### 2.1 Human cells exposed to *B. burgdorferi* strain B31 show altered gene expression

HUVECs and HEK-293 cells are widely used human cell models. As illustrated in Fig 1, we exposed HUVECs and HEK-293 cells to *B. burgdorferi* strain B31 for 72 h to monitor and assess the response, specifically targeting changes in gene expression and concomitant DNA methylation changes.

**Fig 1.**
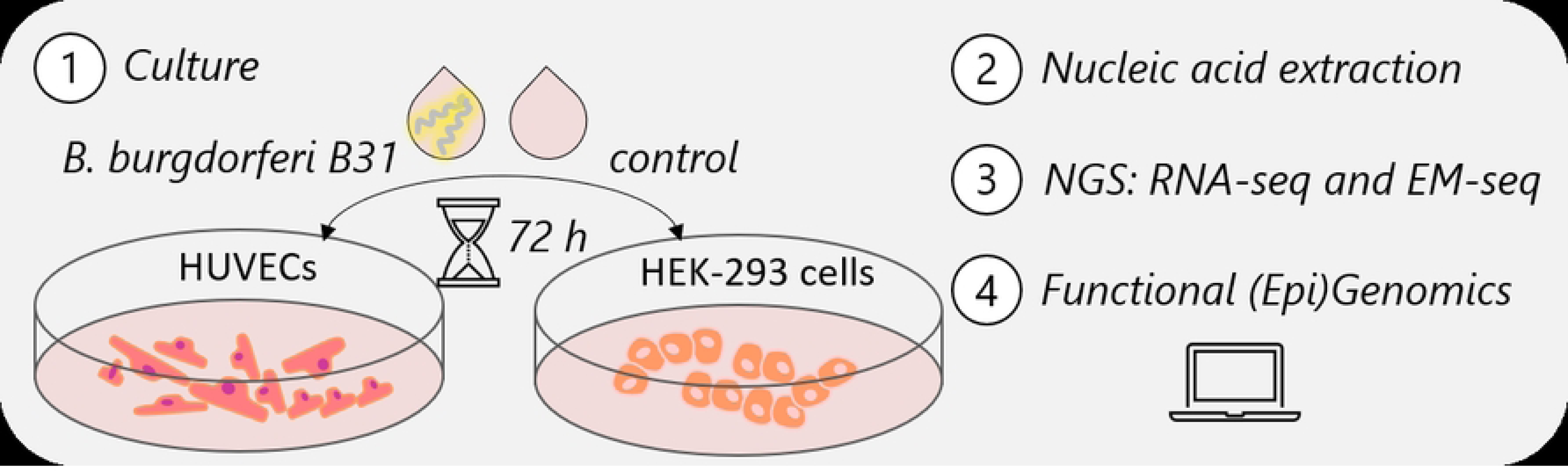
Experimental workflow for human cells exposed to *B. burgdorferi*. HUVEC and HEK-293 cells, were exposed to *B. burgdorferi* B31 for 72 h. RNA and DNA were extracted and subjected to library preparation for next-generation sequencing (RNA-seq and Enzymatic Methyl-seq). Genome-wide epigenomic and transcriptomic data were used for functional enrichment analysis.

The RNA-seq study generated 1.07 billion reads (an average of 89.57 million reads per sample). Mapping efficiency to the reference genome averaged 97.9 % as documented in S1 File. Principal component analysis (PCA) was applied to explore the comprehensive expression data and map the correlation in a quantitative 2D graph, after filtering out low-expression gene loci (Fig 2). We first compared the distribution of unexposed or exposed experimental replicates in each cell model individually. We then also included the two cell types and all treatment conditions in the comparison. For each cell type, there was a clustering along principal component (PC) 1 which explains 24.72 and 22.94% of the variation among unexposed control cells and *B. burgdorferi*-exposed HUVECs and HEK-293 cells, respectively (Figs 2A and 2B). This clustering can be attributed to exposure conditions, which explain the largest variation in the data set. The variation for PC2 could be attributed to the experimental replicates. When HUVECs were compared to HEK-293 cells, including all exposure conditions, 84.74% and 2.16%, of the differences could be attributed to cell type characteristics along PC1 and PC2, respectively (Fig 2C). Thus, the response differs by cell type.

**Fig 2.**
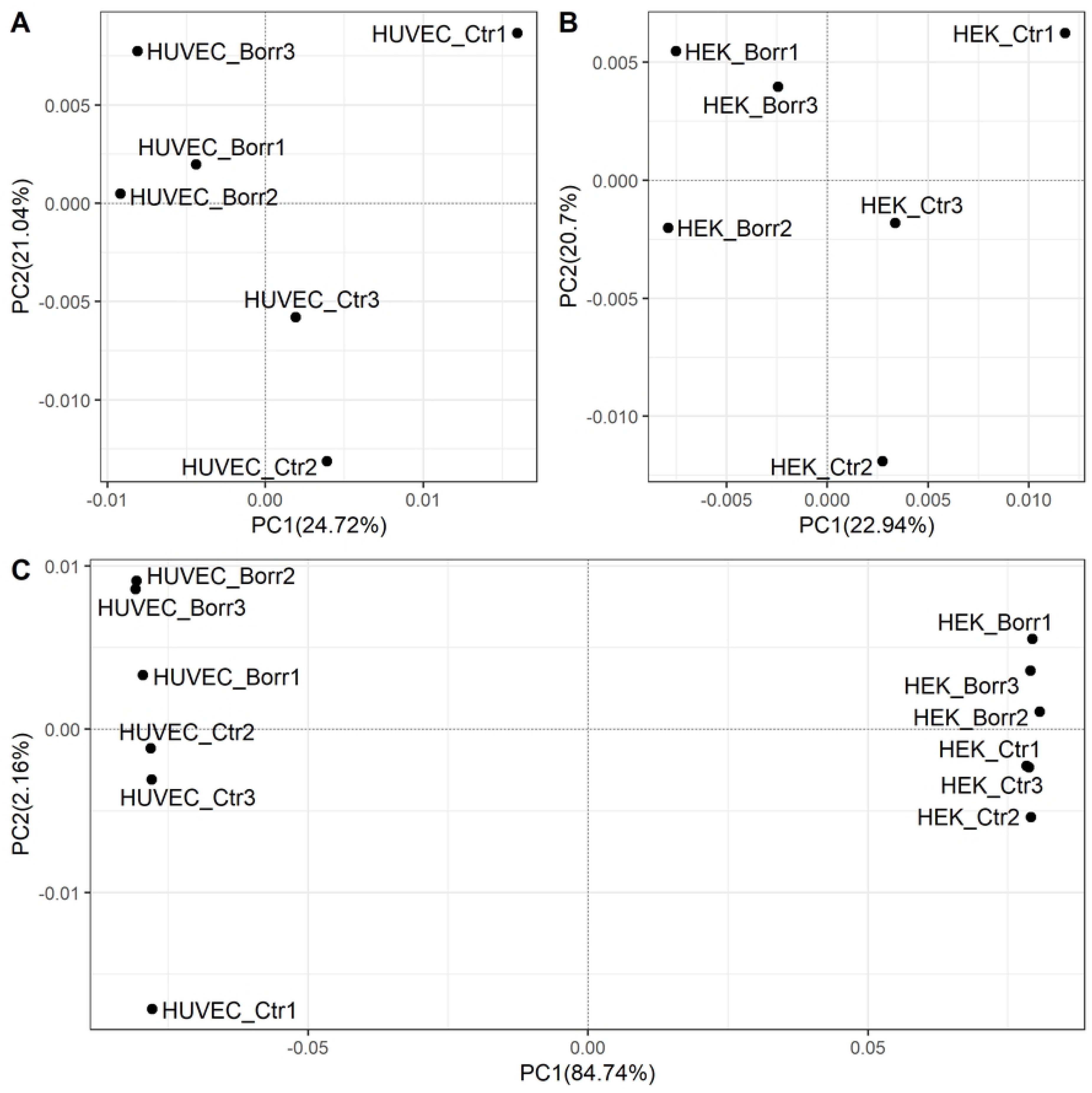
PCA of transcriptome data. Human cell models (A) HUVECs and (B) HEK-293 cells remained unexposed (control; Ctr) or were exposed to *B. burgdorferi* strain B31 for 72 h (Borr). Experiments were performed in triplicate as indicated by the sample names. RNA was subjected to RNA-seq on the NovaSeq 6000 (Illumina). DeSeq2 was used to normalize processed data and to perform gene expression analysis. Data exploration by PCA and visualization was performed in the statistical environment of R and with the R packages EDASeq, ggpubr, and cowplot, with gene loci with low expression filtered out of the data set. Spearman correlation and classical multidimensional scaling were applied. Unexposed (Ctr) and *B. burgdorferi*-exposed (Borr) cells, both for HUVECs and HEK-293 cells, clustered separately along the x-axis with PC 1 explaining the variation between exposure conditions, while PC2 explained the clustering between replicates in each condition. The positively correlated samples are closer together and are located on the same side of the plot. (C) Comparing HUVECs and HEK-293 cells the replicates clustered separately along the X-axis, with PC1 explaining 84.74% of the variation between the cell types. PC2 with 2.16% explained sample correlation clustering. The positively correlated HEK-293 cells conditions clusters are less extended than the cluster for HUVECs.

**Fig 3.**
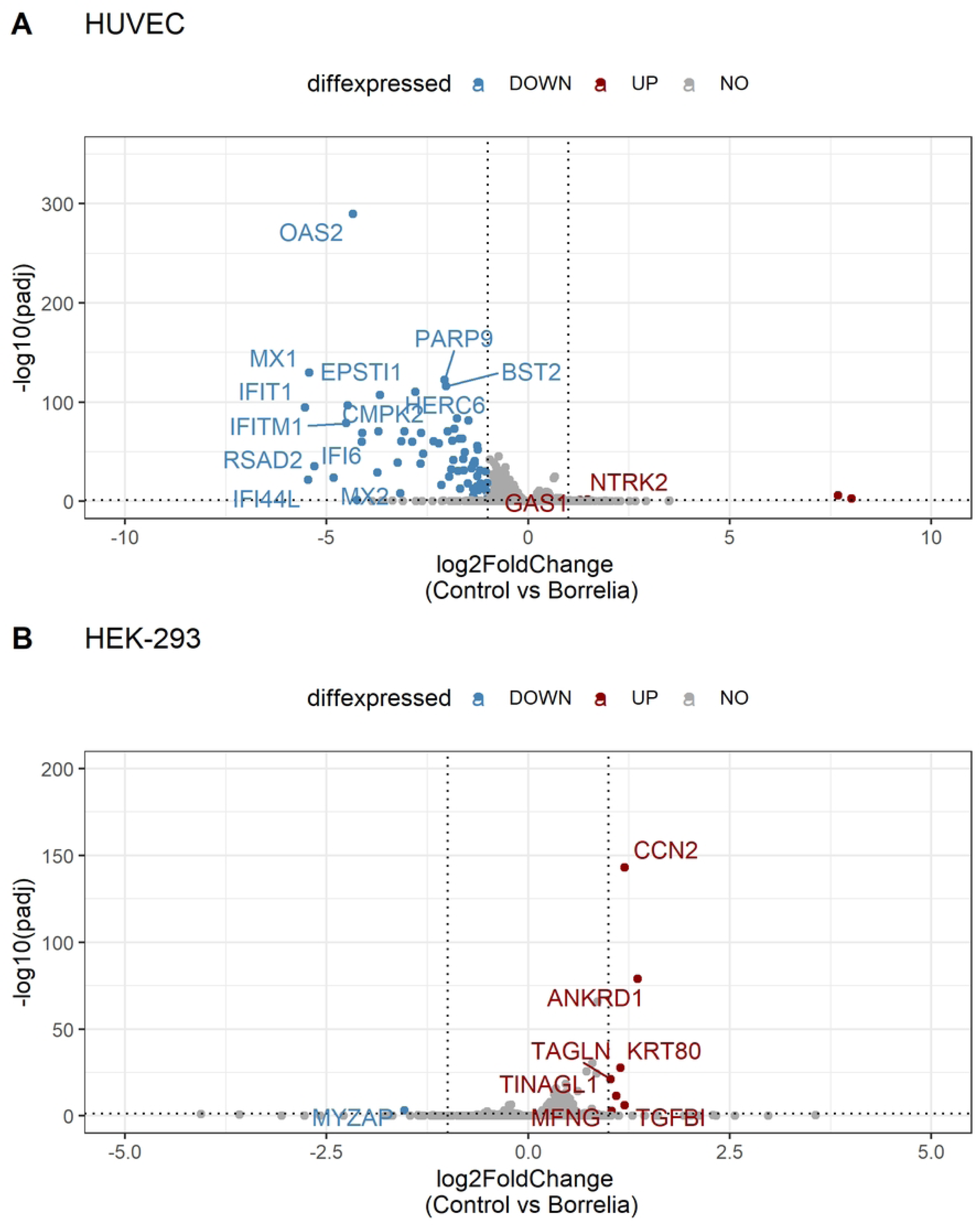
Volcano plot of differential gene expression. Expression data from (A) HUVECs and (B) HEK-293 cells unexposed or exposed to *B. burgdorferi* strain B31 for 72 h were compared. The log2FC on the X-axis, representing the level of change between the two exposure conditions, was plotted against statistical significance in terms of p_adj_ on the Y-axis. The horizontal line marks p_adj_ = 0.05, whereas the vertical lines represent the cut-off of log2FC at - 1 and 1. The blue dots represent the significantly down-regulated genes, while the red dots represent the significantly up-regulated genes in the control-treated cells. The many genes highlighted in gray are outside the cut-offs. HUVECs have considerably more differentially expressed (DE) genes, most of which are downregulated in control cells. In contrast, in HEK-293 cells there are more DE genes upregulated than downregulated in control cells. Graphs were drawn with ggplot2 applying ggrepel in R.

Since the variation difference between cell types was so clear, subsequent analysis treated each cell type separately. Differential expression analysis with DeSeq2 identified the significant changes in the two cell models, HUVECs and HEK-293 cells, in response to *B. burgdorferi* strain B31. The expression data were examined with Relative Log Expression (RLE) plots as quality control to confirm the normalization across samples (S1 Fig).

A total of 26,919 genes in HUVECs (S2 File) were filtered for p-value adjusted for multiple testing using the Benjamini-Hochberg method - *p*_adj_ < 0.05, and 364 differentially expressed (DE) genes were found. Many of these genes were up- or down-regulated less than two-fold, falling in the range of log2 fold change (log2FC) ≤ |1|, but 69 genes had a log2FC ≥ |1|. In HEK-293 cells, a total of 29,367 genes (Supplement File 3) were filtered for *p*_adj_ < 0.05, resulting in 110 DE genes, of which 8 had a log2FC > |1|. Volcano plots (Fig 4) support the findings of the differential expression analysis and demonstrate that in HUVECs there are more genes downregulated in unexposed controls compared to *B. burgdorferi*-exposed cells. In contrast, in HEK-293 cells the majority of significant genes are upregulated in the unexposed control cells compared to *B. burgdorferi*-exposed cells. Heat maps allow visualization of the log2 scaled, mean-centered expression of each of the identified DE genes for each sample replicate in HUVECs (S2 Fig) and HEK-293 cells (S3 Fig). The heat maps also clearly show that the *B. burgdorferi*-exposure produces distinct patterns of gene expression, which is supported by the column dendrograms in the heat maps for the individual cell models. The expression patterns of DE genes also indicate gene clusters in the heat maps supported by the row dendrograms.

**Fig 4.**
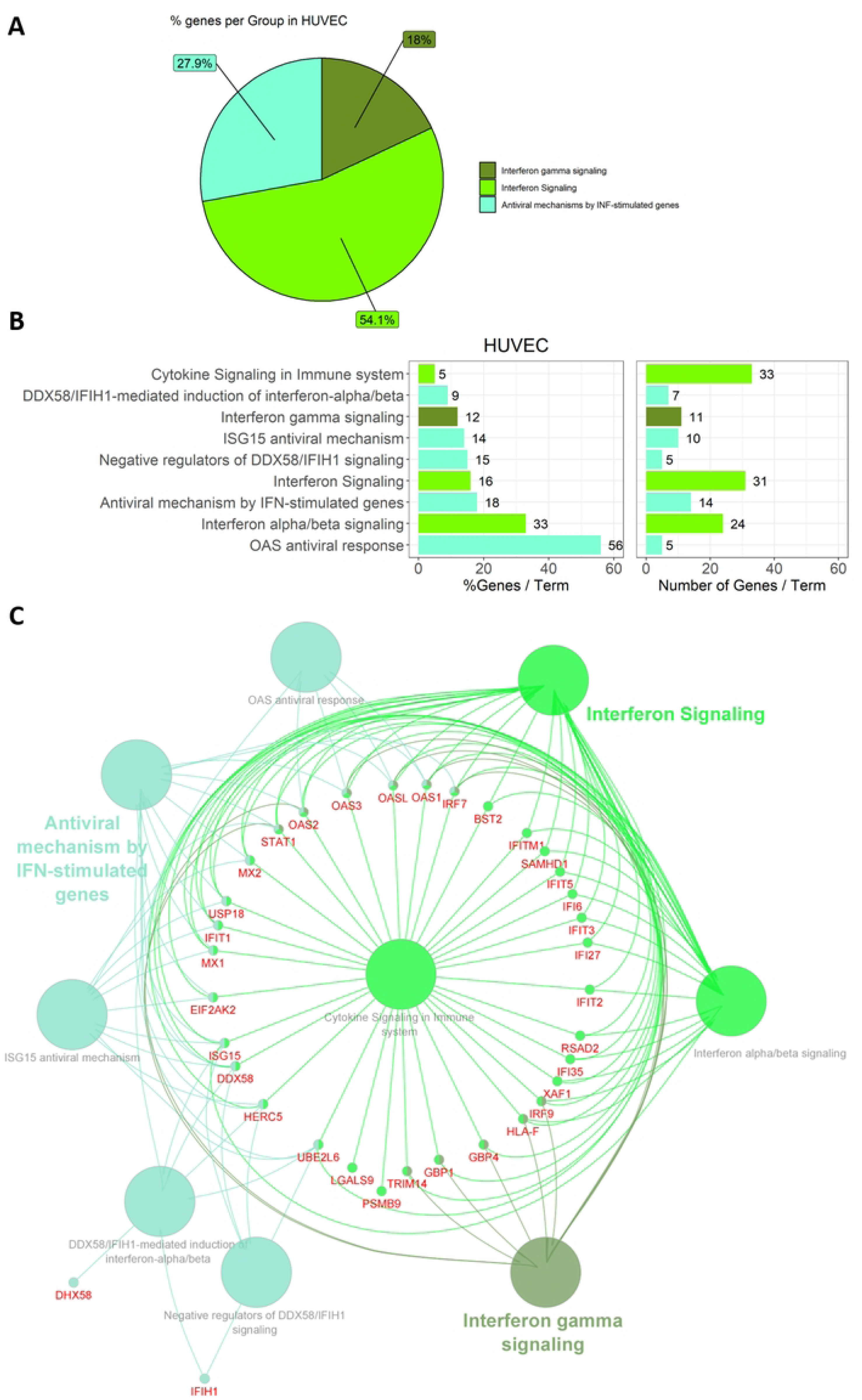
Functional annotation of DE genes in HUVECs. The visualization, in form of pie and bar charts and networks, was enabled with ClueGO in Cytoscape. The subset of 69 differential expressed (DE) genes was imported and assigned to functional pathways of the Reactome database. Enrichment tests were performed and p-values were corrected for multiple testing applying Benjamini-Hochberg. Threshold p < 0.05 was applied for HUVECs in the network of representative pathways. (A) DE genes in HUVECs were assigned to three groups as depicted in the pie chart in three shades of green. (B) Assignment to nine pathway terms was made, with % Gene/Term and Number of Genes/Term mapped separately. (C) The generated network shows the groups (colored), pathway terms (grey), and shared genes (red) of the functional analysis.

### 2.2 HUVECs respond by upregulating host defense genes

The identified DE genes were subjected to functional analysis with Cytoscape App ClueGO using Reactome pathways for enrichment analysis and visualization in networks. Representative pathways with mapped genes, and the percentage involvement in the identified groups and terms, are depicted for HUVECs in Fig 4 and HEK-293 cells in Fig 5.

**Fig 5.**
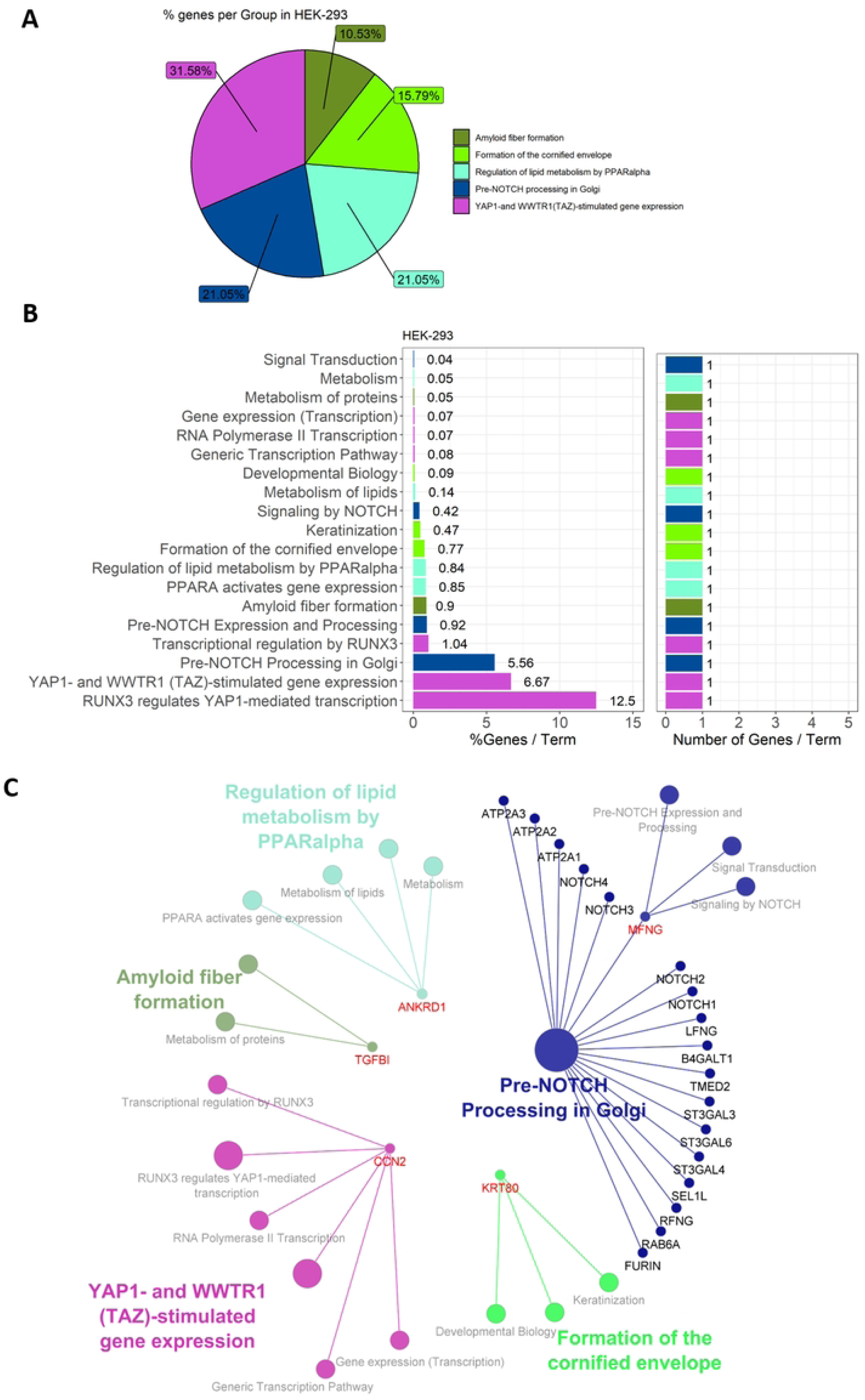
Functional annotation of DE genes in HEK-293 cells. The visualization in form of pie and bar charts and networks was enabled with ClueGO in Cytoscape. The subset of 8 differential expressed (DE) genes has been imported and assigned to functional pathways of the Reactome database. Enrichment tests were performed and p-values were corrected for multiple testing applying Benjamini-Hochberg. Here all potential representative pathways are depicted, while only “Pre-NOTCH Processing in Golgi” is significant. (A) DE genes in HEK-293 cells were assigned to five groups as depicted in the pie chart. (B) Assignment to 19 pathway terms was made. with % Gene/Term and Number of Genes/Term mapped separately. (C) The generated network shows the groups (color-coded), pathway terms (grey), and contributing genes (red) of the functional analysis.

Four of the 69 significantly DE genes were not annotated. Of the remaining 65, 53 (81.5%) were recognized and included in the functional annotation. These genes were associated with 9 representative terms and pathways after p-value significance selection criteria (Fig 4B). They were divided into three groups: "Interferon-gamma signaling", "Interferon signaling", and "Antiviral mechanisms by INF-stimulated genes"; "Interferon signaling" was the group with the highest percent of genes per group (Fig 4A). “Interferon alpha/beta signaling” and “OAS antiviral response” were attributed with the highest percent of genes of a pathway/term, while the number of genes per term varied and most genes were found in “Cytokine signaling in immune system” with 33 genes involved in this pathway (Fig 4B). The generated network for HUVEC (Fig 4C) shows the terms as intersection points linked based on a previously determined kappa value. All three groups with assigned pathway terms shared the genes *STAT1*, *OAS1*, *OAS2*, *OAS3*, *OASL*, and *IRF7* as depicted in the circular layout network of Fig 4C. In HUVECs, exposure to *B. burgdorferi* shows a pronounced induction of gene expression, as documented in Fig 6 for a selection of genes.

**Fig 6.**
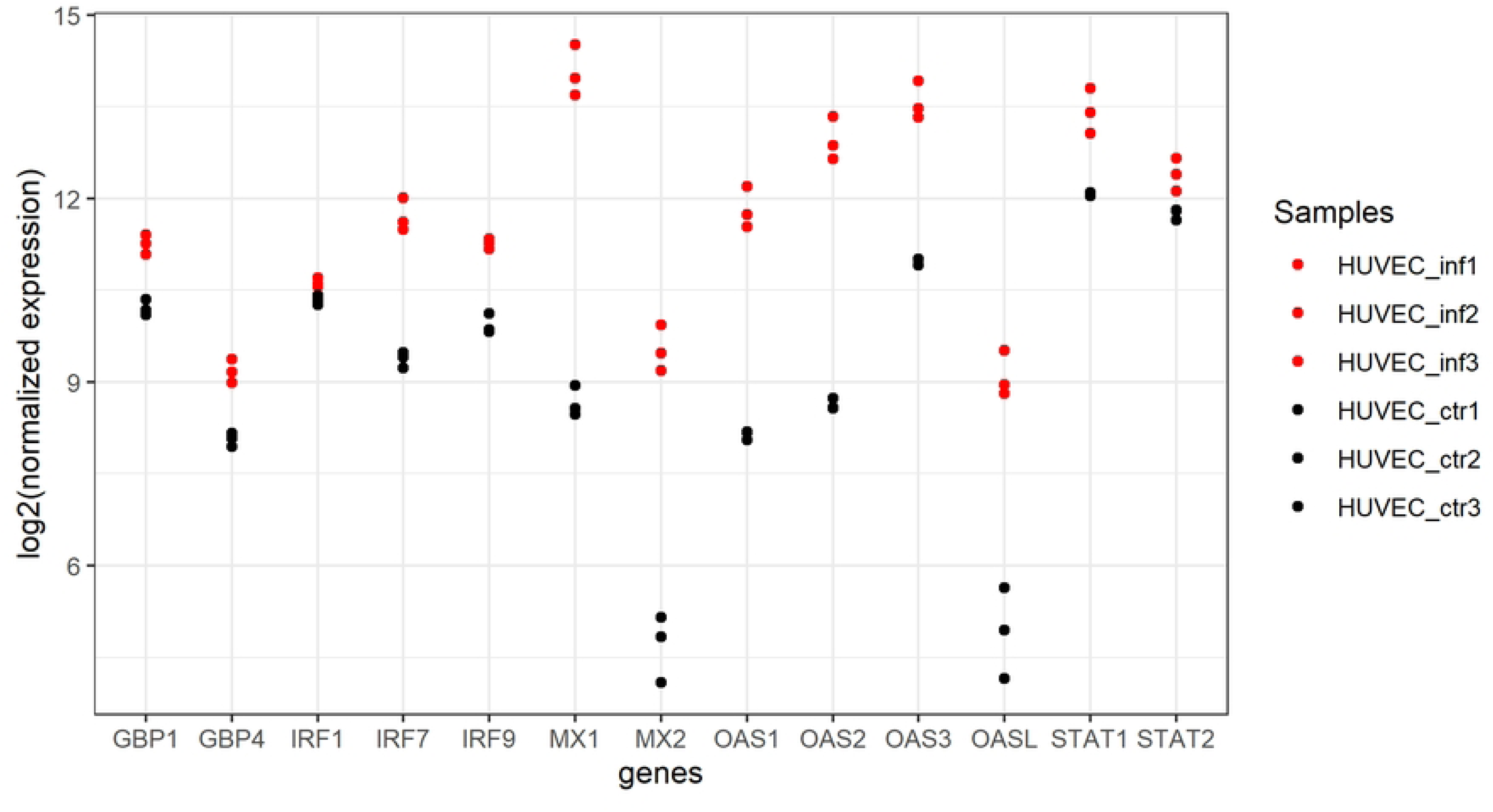
Expression of selected DE genes in HUVECs unexposed or exposed to *B. burgdorferi* strain B31. DE genes passed padj < 0.05, log2FC > |1|. These genes, some of which were also the main players in the functional enrichment analysis used to explain the induced immune response, are clearly shown to be induced in their expression by *B. burgdorferi* exposure. Basic leucine zipper transcription factor (BATF2), C-X-C Motif Chemokine Ligand 11(CXCL11), retinoic acid-inducible gene-I (RIG-I)-receptor (DDX58), guanylate-binding *proteins* (*GBP),* RIG-I-like receptor (IFIH1), Interferon regulatory factors (IRFs), MX proteins that are dynamin-like GTPases, members of the 2’-5’-oligoadenylate synthetase (OAS) protein family, and two Signal Transducer and Activator of Transcription (STAT) are highlighted here from the data set. Genes are plotted against log2(normalized expression) per replicate of the different cell treatments, shown as red dots for *B. burgdorferi*-exposed (Borr) and black dots for uninfected HUVECs (Ctr). The plot was generated with ggplot2 in R.

### 2.3 Gene expression patterns in HEK-293 cells differ distinctly from HUVECs

Of the 8 significantly DE genes in HEK-293 cells, five (62.5%) were functionally annotated and associated with 19 representative terms and pathways after applying general selection criteria (Fig 5). Assignment to groups resulted in 5 groups: "Amyloid fiber formation", "Formation of the cornified envelope", "Regulation of lipid metabolism by PPARalpha", "Pre-NOTCH processing in Golgi", and "YAP1-and WWTR1(TAZ)-stimulated gene expression" (Fig 5A). Because of the short input gene list, every term was associated with only 1 gene, each of which accounted for a different percentage in the signaling pathway (Fig 5B). Applying the Benjamini-Hochberg correction of the pathway analysis only "Pre-NOTCH processing in Golgi" and *MFNG* were returned as significant. Unlike HUVECs, DE genes in HEK-293 cells were mostly downregulated in response to *B. burgdorferi*, except for MYZAP (Fig 7).

**Fig 7.**
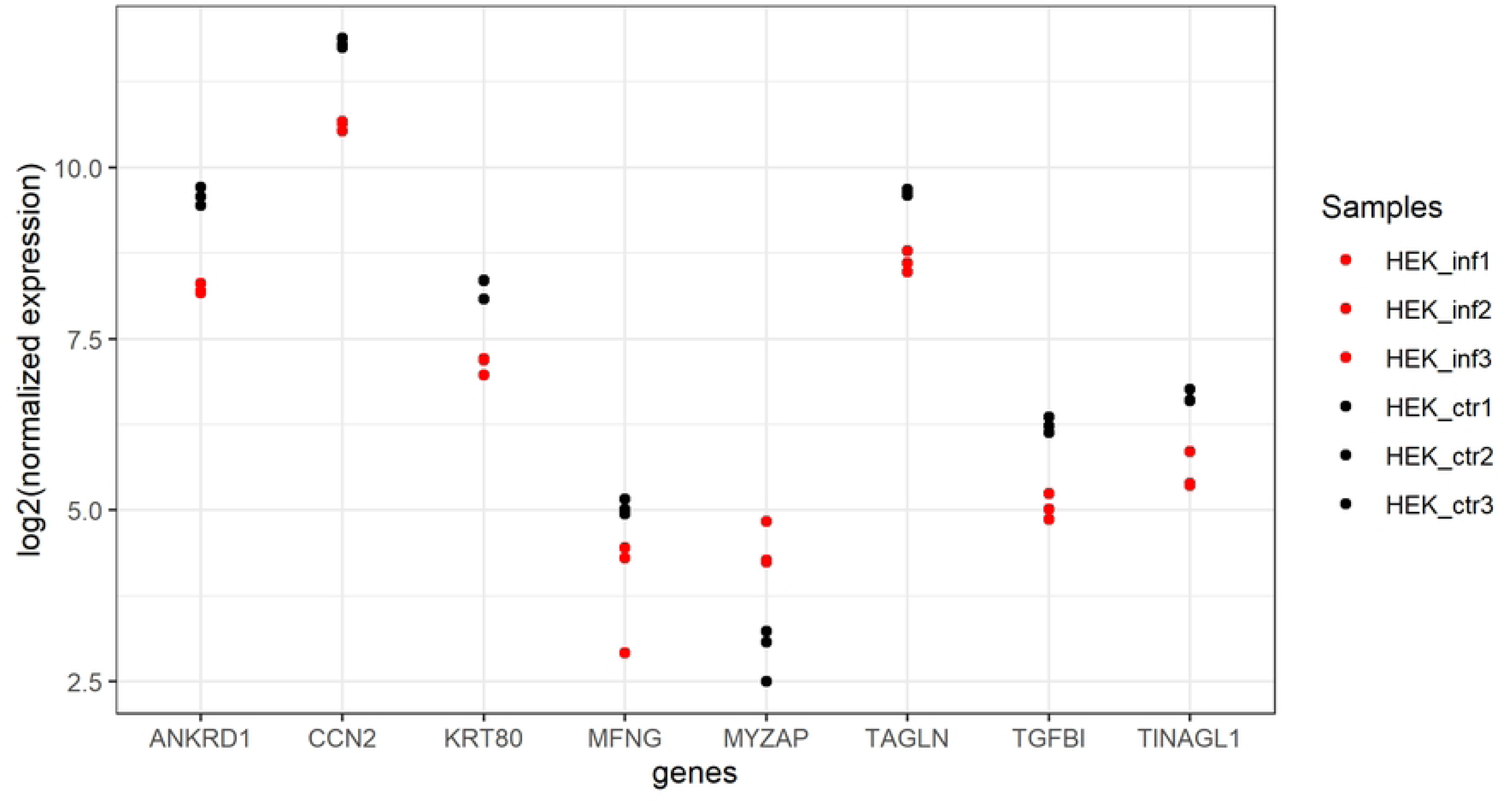
Expression of DE genes in HEK-293 cells unexposed or exposed to *B. burgdorferi* strain B31. DE genes passed padj < 0.05. log2FC > |1|. Most DE genes in HEK-293 cells are downregulated upon *B. burgdorferi*-exposure. The 8 DE genes are coding for the following proteins: Ankyrin Repeat Domain 1 (ANKRD1), Cellular Communication Network Factor 2 (CCN2), Keratin 80 (KRT80), O-Fucosylpeptide 3-Beta-N-Acetylglucosaminyltransferase or manic fringe (MFNG), Myocardial Zonula Adherens Protein (MYZAP), Transgelin (TAGLN), Transforming Growth Factor Beta 1 (TGFB1), and Tubulointerstitial Nephritis Antigen Like 1 (TINAGL1).

### 2.4 **Transcriptional changes are not reflected by methylome changes**

Both cell models exposed to *B. burgdorferi strain B31* were examined for changes in DNA methylation. A total of 5.7 billion reads were obtained from the EM-seq. Reads were aligned (directionally) to the human genome (hg19), and pUC19 and lambda genome using the Dragen methylation pipeline on Illumina BaseSpace. Validation of conversion for methylated spike-in control pUC19 revealed an average CpG methylation of 97.73%, ranging from 95.6 to 98.9%.

Unmethylated spike-in control lambda showed an average CpG methylation of 0.66% with 2 clear outliers of 1.92% and 3.44% CpG methylation (Supplement File 4). These results are in accordance with the quality controls and confirm that the method worked. An average of 215.47 million reads per sample were used for alignment to the human reference genome resulting in an average mapping efficiency of 45.23%. The mapping efficiency to the human genome was relatively low in this study, although FastQC was considered to be good; low mapping efficiency has also been reported for other genomes using Bismark. The conversion metrics for the human genome revealed an average CpG methylation of 69.73% with methylation almost exclusively associated with CpGs and not CHG or CHH. Histograms of % CpG methylation in the forward and reverse strands were comparable for all samples and showed either high or low percent methylation per base (Fig 8). Pearson pairwise correlation analysis between the percent methylation profiles across all samples of HUVECs and HEK-293 cells showed a strong sample correlation independent of the exposure status (Fig 8). This high correlation of the samples indicates that there were hardly any measurable differences due to *Borrelia* exposure. Supposedly due to the primary cell origin, the strong correlation values for HUVECs are slightly lower compared to HEK-293 in all modified settings for differential methylation analysis. CpG coverage was plotted for both cell models to ascertain the quality of reads from EM-seq (Fig 9). A selected CpG coverage of 5 or 10X suggests that the coverage is related to true CpG and not a technical artifact.

**Fig 8.**
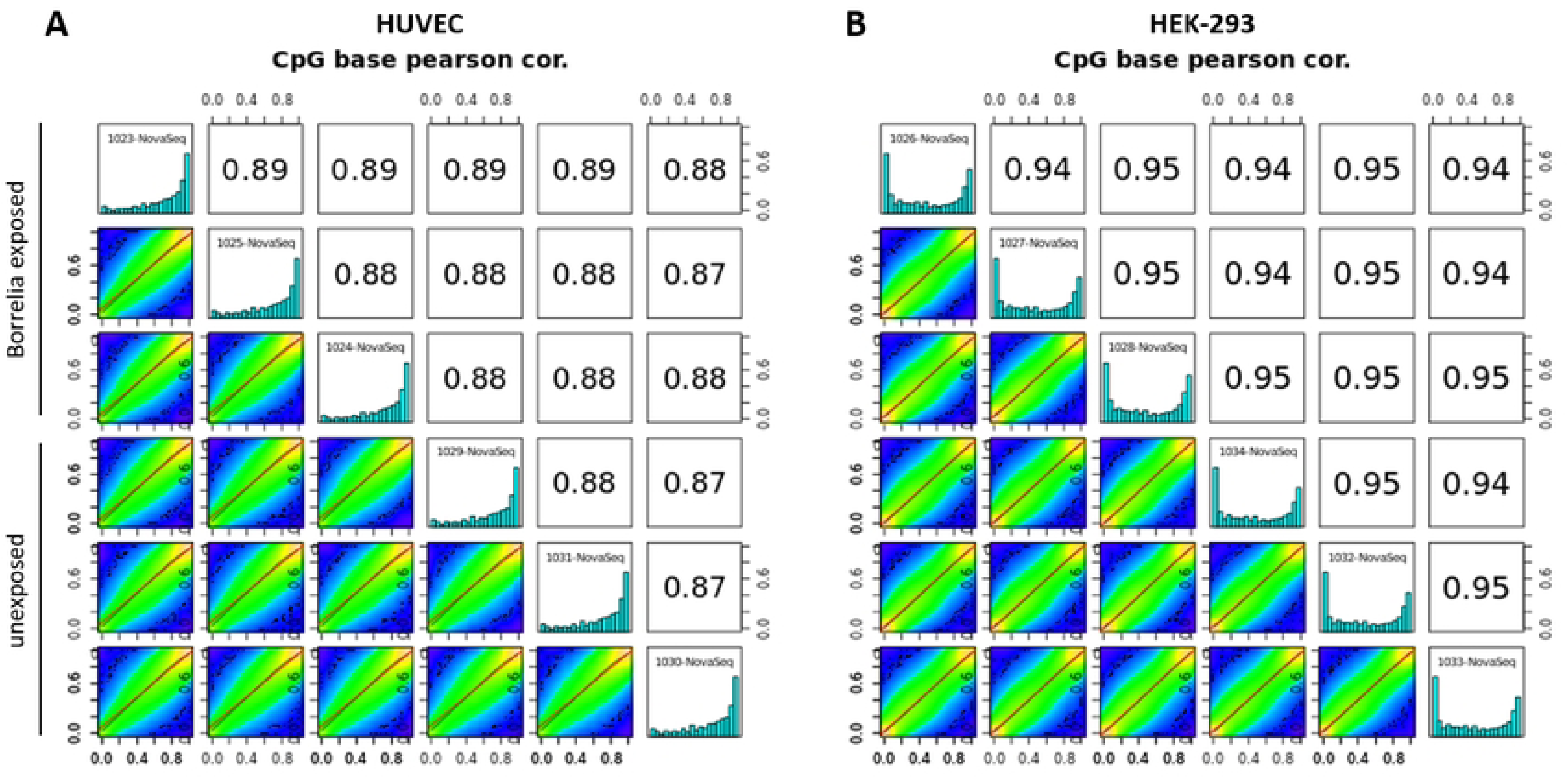
Correlation of methylation across samples. Correlation plots for (A) HUVECs and (B) HEK-293 cells were generated with MethylKit on Illumina Basespace. Histograms displaying percent methylation per cytosine base for unexposed and *B. burgdorferi*-exposed replicates are shown diagonally for each cell model. Numbers in the top right panels indicate Pearson correlation values of the pairwise comparison. The panels in the lower left are scatter plots of percent methylation for each sample pairing. The settings for sample comparison of unexposed and *B. burgdorferi*-exposed cells of the figures shown here were a 20% methylation difference, minimum CpG coverage of 10x, and q-value of 0.05 for both HUVECs and HEK-293 cells.

**Fig 9.**
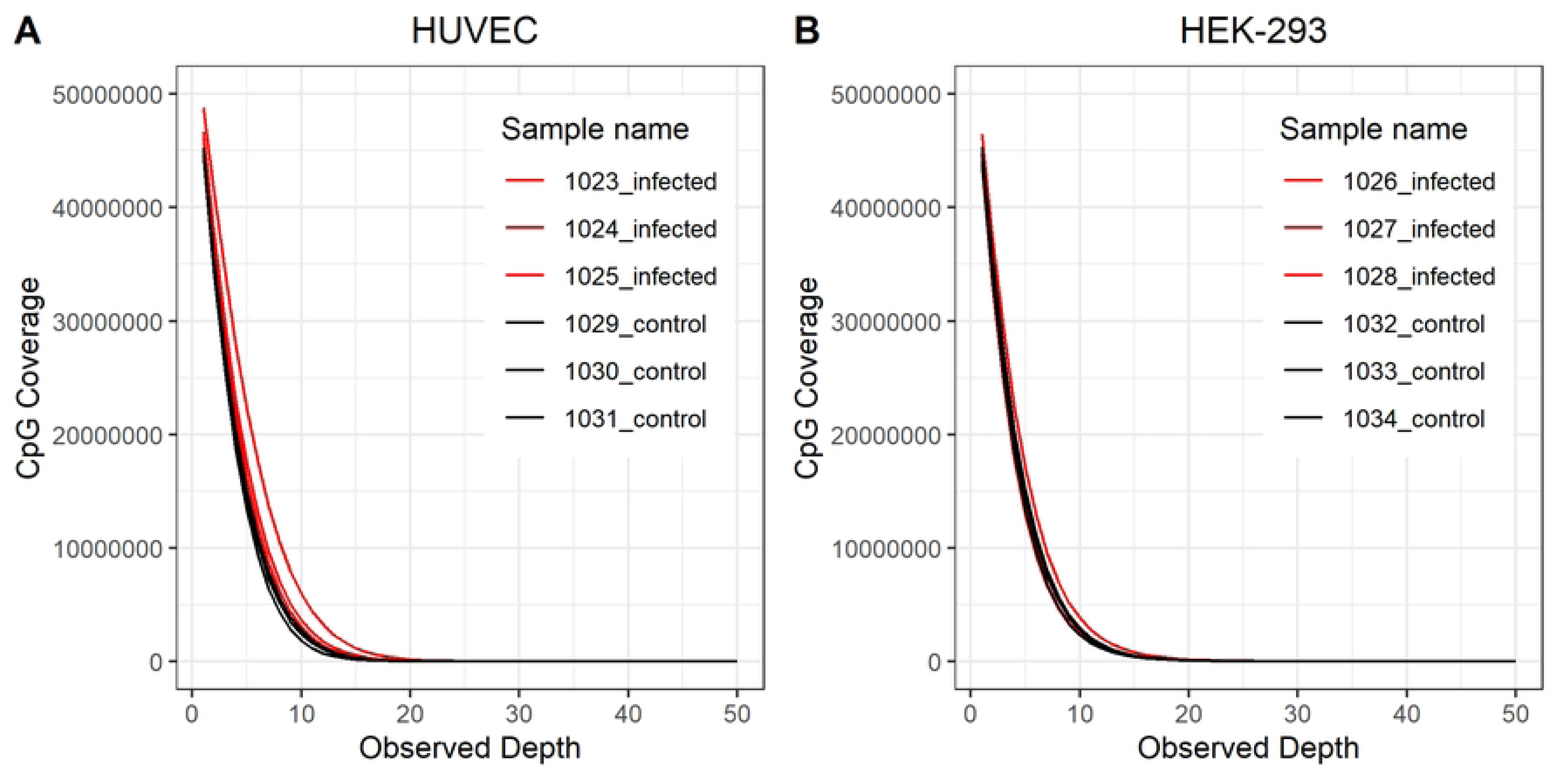
Observed coverage of CpGs after EM-seq in unexposed and *B. burgdorferi*-exposed human cells. DNA was sheared to 300 bp using the sonicator (E210, Covaris) and used as input for EM-seq on the Novaseq 6000. *As* for quality control of methylome analysis, the CpGs covered in (A) HUVECs and (B) HEK-293 cells unexposed and *B. burgdorferi*-exposed samples were plotted against observed sequencing depth. A chosen CpG coverage of 5X or 10X suggests confidence in real CpG and not a technical artifact.

Differential methylation analysis testing variable settings of percent difference and coverage between comparisons, and applying two different q-values in MethylKit did not reveal any significant methylation differences between unexposed and *B. burgdorferi-*exposed cells, in either cell model (S4 Fig). Thus, transcriptional changes are seen but not associated with DNA methylation.

### 2.5 Changes of non-coding RNAs in response to *Borrelia*-exposed HUVEC

Four genes of the DE genes of HUVECs were not annotated with a gene symbol, but they showed a log2FC that was considered significant between *B. burgdorferi* unexposed and exposed cells. Manual inspection revealed that these gene loci encode a pseudogene (N-ethylmaleimide-sensitive factor pseudogene 1 (NSFP1), ENSG00000260075.1), a novel transcript (long ncRNA, ENSG00000272512.1), a novel protein (ENSG00000285304.1), and a processed transcript (PDCD6-AHRR readthrough (NMD candidate) (PDCD6-AHRR), ENSG00000288622.1). Both the novel transcript and the processed transcript are RNA genes that belong to the long non-coding RNA (lncRNA) class according to GeneCards (www.genecards.org). Transcription of the novel transcript (lncRNA) was doubled upon *B. burgdorferi* exposure, while PDCD6-AHRR was downregulated in exposed HUVECs.

### 2.6 Overlap in differentially expressed genes between this study and other studies of *Borrelia*-exposed cells

An integrative computational approach was used to compare the results of this study with relevant previously published transcriptome data. Fig 10 shows Venn diagrams and a table of overlaps and unique gene expression profiles for the two cell models exposed to *B. burgdorferi B31* in this study, versus other studies of host cell-pathogen interaction.

**Fig 10.**
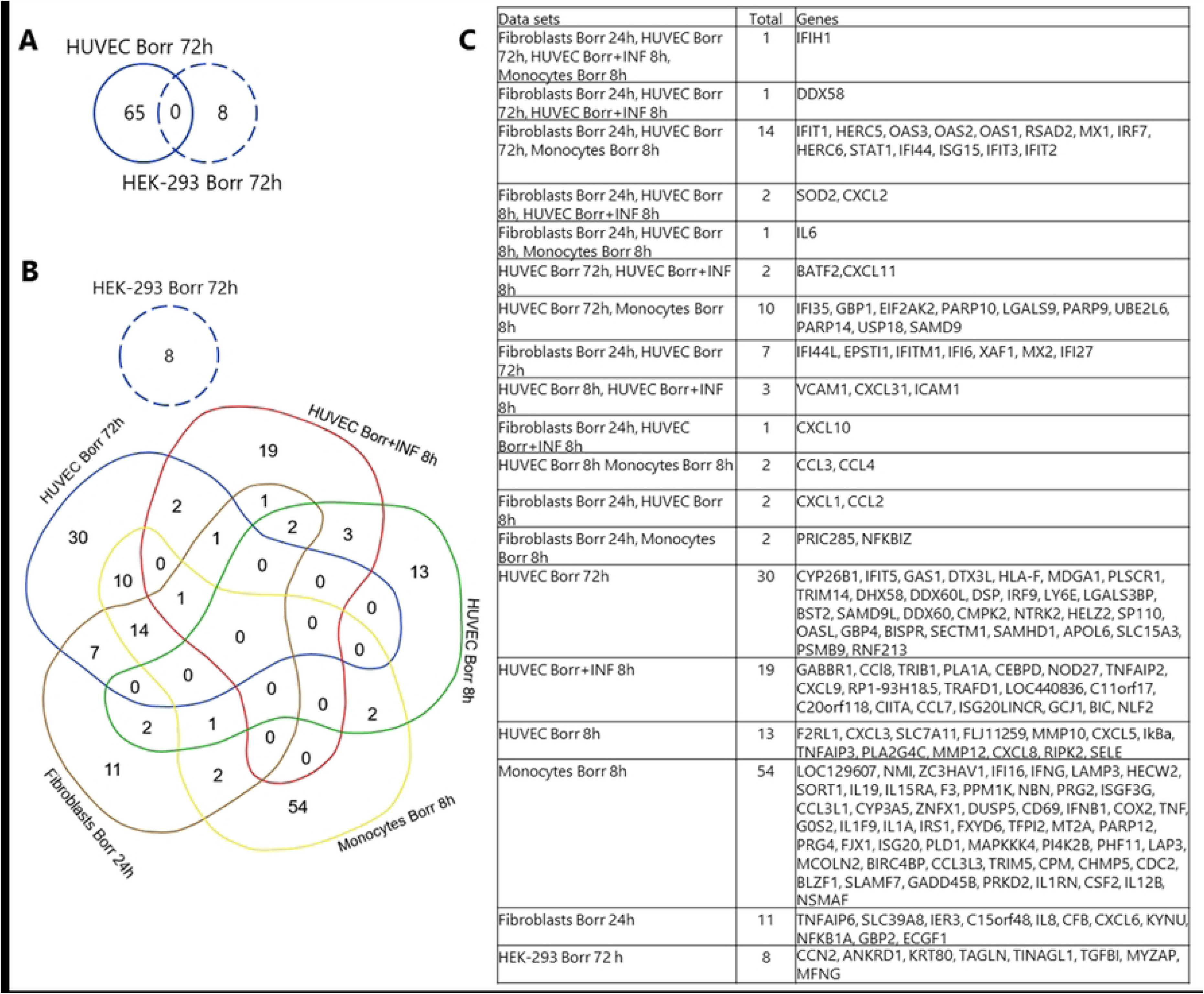
Comparative analysis of gene expression induced by *B. burgdorferi* in different cell types. (A) Differentially expressed (DE) genes identified in this study for HUVEC and HEK-293 cells exposed to *B. burgdorferi* for 72 h were compared and showed no overlap in a Venn diagram. (B) Other transcriptome studies were used for the comparison and served as input for the Venn diagram drawn with the online tool: https://bioinformatics.psb.ugent.be/webtools/Venn/. The data sets are color-coded. Dame *et. al* (2007) examined the effect of *B. burgdorferi* with a MOI 10:1 (green) or *B. burgdorferi* (MOI 10:1) in combination with Interferon-gamma (INF) (red) on HUVECs after 8 h exposure by microarray. Salazar *et al.* (2009) exposed monocytes with lysed and live *B. burgdorferi* for 8 h at various MOIs (1:1,10:1,100:1). From their study, genes exclusively or more intensely up-regulated by live *B. burgdorferi* (yellow) were included in the present comparative study. Meddeb *et al.* (2016) used fibroblasts that were exposed to *B. burgdorferi* and two other pathogenic *Borrelia* strains at a MOI 100:1 for 24 h (brown) subjected to microarray analysis. Consistent DE genes in response to the three pathogenic were considered for this comparative study. The data sets in blue represent the gene set of the present study where HUVECs (solid blue line) or HEK-293 cells (dashed blue line) were exposed to B. burgdorferi for 72 h at a MOI of 50:1. Our HUVEC data set (solid blue) shows no overlap with HUVECs exposed to *B. burgdorferi* for 8 h (green). HUVEC showed some gene overlap with the others studies, while no shared DE genes were found between HEK-293 cells and the other data sets. (C) The table provides the gene names of shared and unique genes from each data set.

Dame *et al.* (2007) studied the effect of *B. burgdorferi* at a MOI 10:1 or *B. burgdorferi* (MOI 10:1) in combination with Interferon gamma (INF) on HUVECs after 8 h exposure by microarray [54]. Dame *et al.* (2007) found that *B. burgdorferi* together with INF-γ induced more genes in HUVECs. Salazar *et al.* (2009) exposed monocytes to lysed and live *B. burgdorferi* for 8 h at various MOIs (1:1,10:1,100:1) [49]. Genes that are upregulated exclusively or more strongly by live *B. burgdorferi* than by dead bacteria were included in the comparative study presented here, as we used live bacteria. In the study by Meddeb *et al.* (2016) fibroblasts were exposed to *B. burgdorferi* and two other pathogenic *Borrelia* strains at a MOI 100:1 for 24 h and subjected to microarray analysis [40]. Genes consistently differentially expressed in response to the three pathogenic *Borrelia* species were considered for this comparative study.

The data show no overlap between DE genes found in HUVEC and HEK-293 cells exposed to *B. burgdorferi* for 72 h (Fig 10A). HUVEC showed some overlap with DE genes derived from transcriptome data from HUVEC, monocytes, and fibroblasts exposed to *B. burgdorferi* for other time periods (Fig 10B). Four genes of our HUVEC data set (solid blue line) overlap with HUVECs exposed to *B. burgdorferi* in combination with INF (red data set in Fig 10B). 25 genes are shared with monocytes exposed for 8 h to *B. burgdorferi* (yellow) and 26 genes are shared with fibroblasts exposed to *B. burgdorferi* for 24 h (brown) and our HUVEC data. In contrast, HEK-293 had no concordant genes with these data sets.

## 3. Discussion

Many questions exist about the interactions between the *Borrelia* pathogen and the host cell that results in Lyme disease. Here, we have focused on representative human cell models to achieve a more detailed understanding of the interactions between *B. burgdorferi* and human cells in a time frame consistent with the early stages of infection when the tick is taking a blood meal from its host. We investigated changes in gene transcription, including non-coding RNA genes, and DNA methylation after 72 h of pathogen exposure. Previous transcriptional studies have used microarray or RNA-seq for blood cells [49, 57], fibroblasts [40, 58], endothelial [54] and neuronal cells [59] and found transcriptional changes after exposure to *B. burgdorferi*. Our study reinforces the occurrence of transcriptional changes induced by *B. burgdorferi*. These occurred to a much greater extent in HUVECs than in HEK-293 cells, and these cell models showed a different spectrum of genes and effects on gene expression.

### 3.1 **Immune response of HUVECs**

The cellular origin differs between the two cell models used in our study. HUVECs are a human primary endothelial cell line. Primary cells are generally considered to be more physiologically relevant than immortalized cell lines. Furthermore, endothelial cells are the first line of defense against infectious agents. Endothelial cells line the body’s blood circulatory system and are immunologically significant due to the production of pro-inflammatory signaling molecules as well as the selective permeability of immune cells into tissues [60, 61]. They are also crucial for nutrient supply and the removal of waste products [62]. Thus, endothelial cells as a cellular barrier between blood and tissues will inevitably come into contact with *Borrelia* sp. during host infection. This occurs either during local infection in the skin or during the spread of infection in the body. Endothelial cells can act as a sensor for the pathogen or provide them with a replicative niche [61].

In our transcriptome study, the cellular immune response in HUVECs was particularly pronounced after *B. burgdorferi* exposure, highlighting its cellular role as an important nexus mediating between the pathogen and host. Pathways enriched for pro-inflammatory cytokines like Interferon-gamma (INF-γ) signaling indicate the linkage of innate and adaptive immune responses that would recruit immune cells to the *B. burgdorferi* infection side *in vivo* [63, 64]. INF-γ was one of the major cytokines found in erythema migrans, one of the early signs of Lyme disease [65, 66]. Dame *et al.* (2007) found that the simultaneous stimulation of endothelial cells with *B. burgdorferi* and INF-γ led to more dramatic transcriptional changes and induction of chemokines [54]. High INF levels have also been found in sera from Lyme disease patients; in patients with late or chronic Lyme disease, these high INF levels indicate a persistent immune response and possible continued infection [54, 65]. In our study, cytokine signaling in the immune system and INF signaling (INF-α, -β, -γ) were induced after 72 h, thus the overall inflammatory picture of the early Lyme disease is reproduced in our study *in vitro* in HUVECs.

Although Dame *et al.* (2007) similarly detected inflammatory response in HUVEC, the majority of genes affected were different compared with our results. Surprisingly, there is no overlap of the *B. burgdorferi*-induced genes in HUVECs incubated for 8 h [54] and those DE genes we found in HUVECs incubated with *B. burgdorferi* for 72 h in this study (Fig 10). In the Dame *et al.* (2007) microarray study, *B. burgdorferi*, IFN-γ, or both showed strong induction of chemoattractant (*CCL7, CCL8, CX3CL1, CXCL10, CXCL2,* and *CXCL9*) and adhesion molecules (*ICAM-1* and *VCAM-1*) in HUVECs after 8 h [54]. Similarly, fibroblasts exposed to three species of *B. burgdorferi sensu lato* for 24 h showed strong induction of chemoattractant molecules [40]. However, for HUVECs induced for 8 h with *B. burgdorferi* in combination with INF-γ [54], there are four common genes (*DDX58*, *IFIH1*, *BATF2*, and *CXCL11*) compared to the present experiments in HUVEC (Fig 10). *DDX58* is a retinoic acid-inducible gene-I (RIG-I)-receptor. *IFIH1* gene encodes MDA5, a RIG-I-like receptor. Both are crucial for the recognition of pathogens in the cytosol and induce IFN-stimulated genes (ISGs), type I IFN, and proinflammatory cytokines [67]. Both genes are also induced by *B. burgdorferi* in fibroblasts [40], and monocytes [49] exposed between 4 and 24 h. Basic leucine zipper transcription factor (BATF2) controls immune cell differentiation and macrophage activation [68]. *CXCL11* is known to be induced by INF-γ, is crucial for activated T-cell recruitment, and has been identified as important for the balance of angiogenesis or angiostasis [69]. These gene products are important connectors between the innate and adaptive immune systems, rely largely on INF signaling, and therefore may play an important role in the early phase of *Borrelia* infection.

A particularly interesting result was that endothelial HUVECs express increased levels of *STAT1*, *OAS1*, *OAS2*, *OAS3*, *OASL*, and *IRF7* after 72 h of *B. burgdorferi* exposure; these genes, are present in all identified pathway groups in the functional analysis. An essential component of the INF signaling pathway via JAK-STAT is the Signal Transducer and Activator of Transcription 1-alpha/beta (STAT1). STAT1 gets either phosphorylated or associated with other factors (STAT2, IRF9), which enables the cytosol-to-nucleus transposition and expression of ISGs that are crucial for response to pathogen stimuli [70]. Among the triggered ISGs are the 2’-5’-oligoadenylate synthetase (*OAS*) protein family, which comprises *OAS1*, *OAS2*, *OAS3*, and *OAS-like* (*OASL*) [71]; all four were found induced by exposure to *B. burgdorferi* in our study. OAS proteins recognize double-stranded RNAs and are thought to have mainly antiviral functions by activating RNase L to eliminate pathogen load through RNA degradation [72]. However, recently *OAS* was also found to be induced by tuberculosis bacteria in macrophages and positively affect the expression of TNF-α and IL-1β to support host defense mechanisms [73]. OASL has also been described as bifunctional with anti- and pro- functionality for viruses and intracellular bacteria [71]. *Borrelia* internalization and cellular processing have been well studied in macrophages [74]. Different pattern recognition receptors in the endosomal compartment recognize cell-invasive pathogens such as bacteria or viruses, and the downstream adaptor proteins induce interferon regulatory factors (IRFs), including *IRF7*, that control and promote the immune response by inducing more ISGs in the nucleus [75, 76]. Another group of ISGs is Guanylate-Binding Proteins, of which *GBP1* and *GBP4* were found significantly expressed in response to *B. burgdorferi* in our study (Fig 6). In addition to their activity against viruses, these proteins have been similarly reported to be active against intracellular bacteria by boosting innate immunity and helping to control tissue damage [77]. GBP1 has a recognition function for cytosolic gram-negative bacteria such as *Salmonella* or *Shigella* and induces the recruitment of additional GBPs and caspase-4 targeting the degradation of the pathogen [78, 79]. Complementing the groups of ISGs found in this study are the proteins MX1 and MX2. They are dynamin-like GTPases, that are induced through the INF signaling pathway via JAK-STAT [80]. *GBP1*, *MX1*, *OAS1*, *OAS2*, *OAS3*, and *IRF7* are among 25 genes shared by monocytes exposed for 8 h to *B. burgdorferi* [49] and DE genes found in this study in HUVECs. ISGs like *IFIT*, *OAS1-3*, *MX1*, *MX2*, or *IRF7* were among the 23 genes shared by fibroblasts exposed for 24 h to *B. burgdorferi* and HUVECs in this study. The different suites of genes found differentially regulated by exposure to *B. burgdorferi* in our study compared to others emphasize the temporal induction of key factors in the immune response, which is crucial for defense against the pathogen in different cell types.

The cellular response we found in this study in HUVECs is primarily aimed at actively eliminating pathogens within the cell. This suggests that *B. burgdorferi* is internalized and is stable in the internal environment long enough to trigger a cellular immune response in HUVECs. *Borrelia* occur both extracellularly and intracellularly [37,81–83]. Karvonen *et al.*, (2021) found some internalized *B. burgdorferi* in human cells after 24 or 72 h exposure despite a relatively high MOI of 200:1 [81]. In the same study, invasion of two non-immune cell lines by *B. burgdorferi* was shown to differ in terms of attachment and bacterial morphology; localization of the bacteria was observed in different cellular compartments, but not the lysosomes, suggesting *B. burgdorferi* has strategies for intracellular survival [81].

### 3.2 **The cellular response of HEK-293 cells suggests ECM remodeling that could promote bacterial survival**

The properties of HEK-293 cells, being derived from a human embryo, are complex [84]. The exact origin of the cells from the embryo is unclear and characteristics of epithelial cells, mesenchymal cells as well as neuronal cells are attributed to these cells [85]. All three of these cell types are also resident in the dermis and are involved in interactions between *Borrelia* and the host so immortalized HEK-293 cells are relevant as a cell model in this study. HEK-293 cells possess a repertoire of receptors for pathogen recognition, such as the plasma-membrane residing Toll-like receptor (TLR) 5, the endosome associated TLR3, and the cytosolic nucleotide-binding oligomerization domain NOD1, but, importantly, lack surface-expressed TLR2 [86–88]. TLR receptors induce an inflammatory signaling cascade through NF-κb or IRF pathway. In monocytes, TLR2 and TLR8 contribute to the effective degradation of the *Borrelia* pathogen in the phagosome [89]. *In vivo*, TLR2 deficiency in mice has been reported to result in persistently elevated *Borrelia* loads in tissues [90] and genetic variation in TLRs corresponds with certain infectious disease risks [91]. Therefore, studying host response to *Borrelia* in cells with some but not other receptors of pathogen recognition could contribute to understanding strategies of bacterial survival within the host cell.

Integrins mediate cellular signaling and communication from the extracellular environment and as such act as recognition molecules for *Borrelia* [37,92,93]. Interaction between *Borrelia* and the host includes binding and degradation of the extracellular matrix (ECM), through induction of metalloproteases [94, 95], induction of Src family kinases, or reorganization of actin filaments [37]. All these mechanisms allow host cell invasion and spread of the pathogen through host tissues. In our study, we found that *TGFB1*, *TINAGL1*, and *CCN2* were downregulated in HEK-293 cells after *B. burgdorferi* exposure. All of these gene products are thought to be responsible for, or contribute to, the binding and organization of the ECM. Transforming Growth Factor Beta 1 (*TGFB1*) encodes a secreted ligand of the TGF-β (transforming growth factor-beta) protein family. As a latent TGF-β binding protein complex, these proteins interact with fibrillin in the ECM of cells [96]. TGF-β signaling contributes to a multitude of cellular processes including extracellular matrix remodeling [97]. Tubulointerstitial nephritis antigen 1 (TINAGL1) is an abundant matricellular renal protein in the basal layer of the ECM [98] and plays a modulatory role through interaction with ECM elements, such as fibronectin and integrins [99]. One member of the Cyr61/CTGF/NOV (CCN) protein family, CCN2 also termed connective tissue growth factor (CTGF), plays a role in the survival of neuronal and other cells [100] and normal production of vascular basement membranes [101]. This matricellular protein interacts with integrins, low-density lipoprotein receptor-related proteins, growth factors, and ECM components [101], thus contributing to ECM function. Down-regulation of these gene products in HEK-293 cells upon *B. burgdorferi* exposure, all with functions involving ECM interaction could favor bacterial survival and invasion. The absence of TLR2 in HEK-293 cells, one of the recognition receptors of *Borrelia*, also suggests that *Borrelia* are unrecognized and survive.

Basic cytokeratin KRT80 is a cytosolic filament that generally has a structural function in epithelia, interacts with cell-adhesive complexes, and its expression is modulated during cell differentiation [102]. KRT80 was downregulated by half in *B. burgdorferi*-exposed cells relative to control cells, which could suggest complex *B. burgdorferi-*induced changes interfering with cellular structure and signaling. The only gene in HEK-293 cells that was induced upon *B. burgdorferi* exposure encodes the Myocardial Zonula Adherens Protein (MYZAP or MYOZAP). In addition to detection in cardiac myocytes [103] and endothelial cells [104], the presence of MYZAP has been demonstrated in adherens junctions in various epithelia, including cancer cells [105]. The presence of MYZAP in cell-cell contact structures in epithelia is thought to be essential for proper scaffold function which also requires actin filaments and actin-binding proteins [105]. It remains to be elucidated what role up-regulation of MYZAP plays in the presence of *B. burgdorferi*.

O-Fucosylpeptide 3-Beta-N-Acetylglucosaminyltransferase or manic fringe (MFNG) is downregulated in *B. burgdorferi*-exposed HEK-293 cells in comparison to unexposed cells. MFNG participates in the pathway “Pre-NOTCH signaling in Golgi”. NOTCH signaling is a highly conserved signaling pathway that contributes to differentiation, proliferation, and apoptosis depending on the cell context [106]. NOTCH receptor precursors (Pre-NOTCH) are post-translationally modified by the addition of O-glucose, O-fucose, and O-N-acetylglucosamine (GlcNAc) by members of the fringe family (lunatic, manic, and radical fringe) in the Golgi membrane [107]. Interfering with the glycosylation machinery alters signal recognition and downstream cellular processes, which might benefit *Borrelia* survival, potentially by suppressing apoptosis.

### 3.3 **Ticks as a potential instigator of epigenetic changes**

We wanted to determine whether the observed transcriptional changes induced by *B. burgdorferi* were due to underlying DNA methylation changes, but this was not the case. However, this does not allow us to conclude about DNA methylation changes over longer (or shorter) time periods. It is possible that a 72-h exposure was not long enough to induce *de novo* DNA methylation in the human cells; prolonged co-culture of cells with *B. burgdorferi*, representing the persistence of the host infection, might induce epigenetic changes. The contribution of other epigenetic mechanisms like histone modifications or non-coding RNAs (ncRNAs) remains to be explored.

However, in our HUVEC dataset, we found potential evidence of epigenetic modification by identifying pseudogene NSFP1, a novel transcript (long ncRNA (lncRNA)), and PDCD6-AHRR (lncRNA) as significantly differentially expressed. The as-yet-unidentified function of pseudogene NSFP1 could be in the form of a protein or ncRNA, or it could involve regulation of the 3D structure of DNA and its accessibility [108]. These gene products, in the form of long or short ncRNAs, could impact the transcriptional machinery and gene function with a pleiotropic effect on cellular signaling [109]. The deregulation of lncRNAs and interaction with microRNAs has been described in the literature to occur in the context of infectious diseases [110]. The influence of ncRNAs on host defense signaling has been described in the context of bacterial and viral pathogens, some of which are beneficial for pathogen survival in the host [110]. The function of these ncRNAs in the cell and consequences of altered expression during *B. burgdorferi* infection remains to be further explored.

Additionally, genome-wide mapping of the *B*. *burgdorferi* transcriptome identified numerous putative ncRNAs [111]. These extracellular dsRNA or intracellular dsRNA could potentially trigger “viral mimicry” signaling through INF, also potentially inducing epigenetic changes in the host cell.

The environment is the classic trigger of epigenetic changes. But the role of ticks in inducing epigenetic changes in their host remains largely unexplored. In nature, ticks most often transmit multiple pathogens and opportunistic infections can occur in response to immune suppression from tick saliva and the resulting disease. Each of these pathogens interacts with each other as well as with the host in a battle to establish infection and find their niche to survive and spread. Although the tick vector was not considered in our study, in addition to the tick-borne pathogens tick feeding injects tick saliva into the host, and tick saliva is highly complex [112]. Biologically active molecules in tick saliva possess anticoagulant, antiplatelet, vasodilator, anti-inflammatory, and immunomodulatory effects [22]; these components have the potential to modulate the host epigenome. A particularly interesting component of tick saliva is the tick histone protein H4, a protein identical to the human histone H4 [38]. Best known for its structural role in chromatin, tick histone H4 also has an antimicrobial effect on skin commensal bacteria, but not on *B. burgdorferi* [38]. Substitution of humans with tick histone 4 suggests one avenue for modulation of the host epigenetic machinery. Similarly, the heat shock proteins found in tick saliva [113], which are immunogenic proteins, can also be epigenetic modulators.

## 4. Conclusion

In this study, we documented a cellular immune response in our endothelial cell model, HUVECs, in response to exposure to *B. burgdorferi*. This response persisted over 72 h and is common to other cell types and other time points. Our epithelial cell model, HEK-293 cells, demonstrated changes in the expression of genes encoding ECM-resident or interacting proteins. Changes in DNA methylation were not detectable in these cells, however, ncRNAs were deregulated in response to infection suggesting that other epigenetic mechanisms may underlie these changes in gene expression. These findings are summarized in Fig 11.

**Fig 11.**
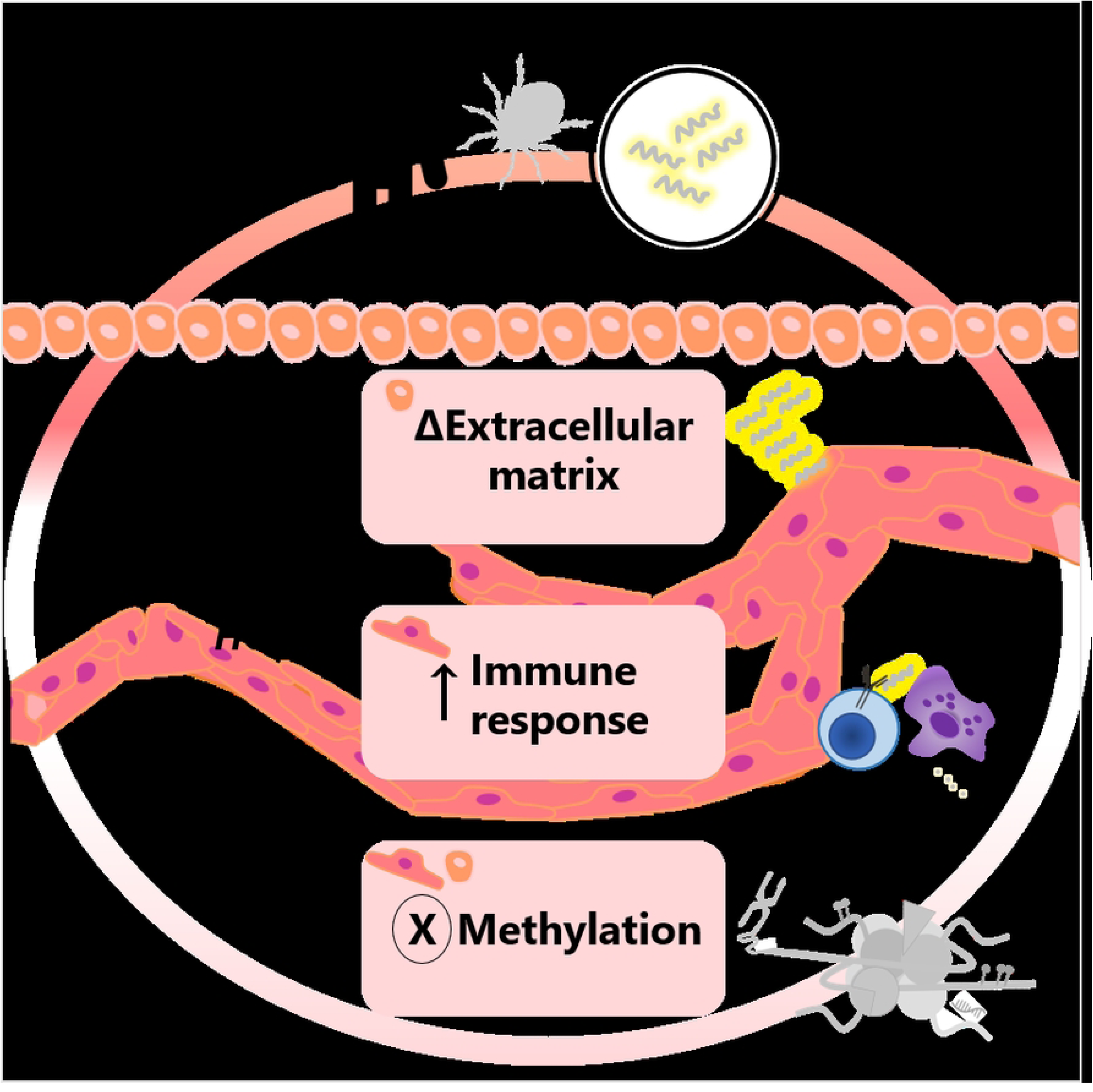
Summary of findings in human cells exposed to *B. burgdorferi*.

## 5. Material and Methods

### 5.1 **Human cell culture**

HUVECs and HEK-293 cells were both originally purchased from American Type Culture Collection and kindly provided by Dr. Gilles Robichaud (Université de Moncton, New Brunswick, Canada). HUVECs (ATCC, PCS-100-010) required optimized endothelial cell growth media EGM^TM^ BulletKit^TM^ (Lonza, CC-3124), which contains EBM^TM^ Basal Medium complemented with 2% v/v FBS, 0.4% v/v bovine brain extract, each 0.1% v/v hydrocortisone, human epidermal growth factor, ascorbic acid, and gentamicin sulfate amphotericin-B (GA-1000). HEK-293 cells (ATCC, CRL-1573) were grown in DMEM media (Lonza, 12-733F) supplemented with 10% v/v heat-inactivated FBS (Rockland, FBS.02-0500), each 1% v/v L-glutamine (Lonza, 17-605E), sodium pyruvate (Lonza, 13-115E), and penicillin/streptomycin (Sigma-Aldrich, P4333). In our study, human and bacterial cell cultures required synchronization. Cells were initially seeded into 75 cm^2^ flasks (Celltreat, 229341) and maintained until 80% confluency with regular media changes every second day before they were counted with trypan blue staining to distinguish living and dead cells (Thermo Fisher Scientific, 15250-061), and seeded into well-plates. Cell growth was monitored with an inverted microscope (Motic AE31). An aliquot of both cell lines was subjected to mycoplasma testing (e-Myco™ VALiD Mycoplasma PCR Detection Kit, Froggabio, 25239) and confirmed negative.

### 5.2 Borrelia burgdorferi culture

*Borrelia burgdorferi* strain B31 (B31 Johnson *et al.* emend. Baranton *et al.*) obtained from American Type Culture Collection (ATCC, 35210) was grown in an optimized nutrient-rich Barbour-Stoenner-Kelly medium including 6% rabbit serum (BSK-H, Darlynn biologicals, BB83-500), supplemented with antibiotics (Sigma-Aldrich) at the final concentration of 50 µg/ml Rifampicin (R3501), 20 µg/ml Phosphomycin (P5396), and 2.5 µg/ml Amphotericin B (A9528) in glass culture tubes (according to [114]). Growth and motility were monitored with a Leitz phase-contrast microscope, and bacteria were counted with a hemacytometer at 250X magnification. All exposure experiments were performed with *B. burgdorferi* strain B31 in an early passage (4 or 5 from original culture) in their exponential growth phase at a concentration of 2 to 4x10^7^ cells/ml, which were grown in a liquid medium at 34°C without shaking, aeration or CO_2_.

### 5.3 Human cell exposure experiment

As shown in Fig 1, starting material for the exposure experiments were HUVECs in passage 3 and HEK-293 cells in passage 5 seeded at a density of 4000 cells/cm^2^ into 6-well tissue culture plates (Celltreat, 229105) in antibiotic-containing media. The three independent replicates of cell exposure experiments were performed with cells of the same passage. At a density of 1.2x10^5^, human cells were exposed to *B. burgdorferi* strain B31 with a multiplicity of infection (MOI) of bacteria cells to human cells of 50:1. Much like natural infections, the *Borrelia* dose per cell, defined as MOI, used in studies ranges from 1 to 1000 in different studies [36–38,40,49,81,115,116]. To our knowledge, there is also no quantitatively validated study on how many *Borrelia* infect a human cell. Presumably, not all tick-borne pathogens transfer from the tick to the host. A 2020 Swedish study quantified typeable *Borrelia* species using the LUX real-time PCR assay and found between 2.0 × 10^0^ to 7.0 × 10^5^ *Borrelia* cells per tick collected from migratory birds [117]. Even if all are transmitted, once in the host, there are multiple host cells and a complex extracellular environment with which the *Borrelia* interact. Consistent with previous studies, we found that human cells were not apoptotic at MOIs of 300 or 600 bacteria per cell at 24 and 48 h (data not shown).

The adherent mammalian cells were washed with PBS (Thermo Fisher Scientific, 14190144) before the addition of resuspended bacteria. *Borrelia burgdorferi* strain B31 cultures were counted, centrifuged at 4500 x g for 10 min and 6x10^6^ bacteria were used per well. The pelleted bacteria were resuspended in human cell growth medium (EBM for HUVECs, DMEM for HEK-293 cells) without antibiotics (Gentamicin-Amphotericin or Penicillin/Streptomycin). In parallel, the cells were incubated with growth media only (without antibiotics) as a control. All plates were incubated for 72 h at constant environmental conditions of 37°C, 5% CO_2_, and humidified air. Human cell growth and morphology were monitored microscopically. After the 72-h exposure, the wells of bacterial-treated or control, for each cell type were washed with PBS and trypsinized and pairs of wells from the same treatment were combined in a 15 ml reaction tube to yield approximately 2x10^6^ cells for nucleic acid extraction. Cells were pelleted by centrifugation at 300 x g for 5 min, the supernatant was removed and cell pellets were kept at -80°C until further use.

### 5.4 Nucleic acid extraction and assessment

All replicates were processed following the manufacturer’s instructions. RNA extraction was performed with the RNeasy Mini Kit (74104, Qiagen) including DNase I on-column digestion (1010395, Qiagen) and elution in 50 µl RNase-free water. The quality and quantity of RNA were initially assessed with a Nanodrop 1000 (Thermo Fisher Scientific) and denaturing RNA-gel. RNA quality was additionally assessed for integrity and size distribution using the Tapestation RNA assay (Agilent, 5067-5576 and 5067-5577), while RNA concentration was determined using the RNA HS assay from Qubit (Thermo Fisher, Life Technologies, Q32852). Samples with a RIN ≥ 9 were used for library preparation.

DNA extraction was performed with DNeasy Blood and Tissue Kit (69506, Qiagen) including an RNase A treatment (R6513, Sigma-Aldrich), and extracted in 100 µl in Tris-HCl pH 8 with 0.1 mM EDTA. DNA concentration was initially estimated spectrophotometrically with a Nanodrop 1000. Next, the quality of the DNA was assessed using the Genomic DNA assay on the Tapestation (Agilent, 5067-5365 and 5067-5366) for integrity and size distribution. Concentration was determined with the dsDNA BR assay from Qubit (Thermo Fisher, Life Technologies, Q32850). Samples with a DIN ≥ 8 were used for library preparation.

### 5.4 Library preparation

#### RNA-seq

The library for RNA-seq was prepared with 200 ng RNA input and the NEBNext Ultra II Directional RNA Library Prep Kit for Illumina (New England BioLabs (NEB), E7760L) at the Atlantic Cancer Research Institute, Moncton, New Brunswick, Canada. First, mRNA was enriched using the NEBNext Poly (A) mRNA Magnetic Isolation Module (NEB, E7490L) and following the manufacturer’s instructions. Briefly, after isolation, samples were fragmented for 15 min at 94°C targeting an insert size of 200 bp. Next, the first and the second cDNA strand were synthesized, followed by a clean-up with Ampure XP beads (Beckman Coulter, A63881). Then the samples were end-prepped and the adaptor was ligated. The adaptor was diluted following the manufacturer’s instructions. The adaptor-ligated samples were purified with Ampure XP beads before amplification. A different Unique Dual Index Primer Pair (NEB, E6440S) was added to each sample and 11 cycles were used for the PCR enrichment. Purification using Ampure XP beads was used to obtain the final library. The quality and quantity of the library were assessed with the D1000 assay on the Tapestation (Agilent, 5067-5582 and 5067-5583) and the dsDNA HS assay from Qubit (Thermo Fisher, Life Technologies, Q32851).

#### Enzymatic Methyl-seq (EM-seq)

The library for EM-seq was prepared using 200 ng DNA input and the NEBNext Enzymatic Methyl-seq Kit (NEB, E7120S) following the manufacturer’s instructions for a standard insert (370-420 bp). First, CpG methylated pUC19 control and unmethylated lambda control were diluted 100 times in TE Buffer pH 8.0 (Thermo Fisher, Life Technologies, 12090015) for final concentrations of 0.001 ng/µl and 0.02 ng/µl respectively. Then DNA samples, and diluted pUC19 and lambda controls were fragmented using the *Covaris E210* Focused Ultrasonicator (Covaris) for a target insert size of 350 bp. After shearing, 1 µl each of pUC19 and lambda controls were added to each DNA sample. Samples were end-repaired, dA-tailed and the adaptor was ligated. The adaptor-ligated samples were purified with NEBNext Sample Purification Beads. EM-seq conversion reaction was performed following the manufacturer’s instructions, starting with the oxidation of 5-Methylcytosines and 5-Hydroxymethylcytosines. DNA-converted samples were purified before denaturation. For denaturation, sodium hydroxide was used and immediately followed by the deamination of cytosines. The deaminated samples were purified before being amplified. For the PCR enrichment, a different EM-Seq Index Primer (NEB, E7120S) was added to each sample and 4 cycles were used. A final purification using NEBNext Sample Purification Beads was performed to obtain the final library. Size distribution of libraries was determined with the D1000 assay on the Tapestation (Agilent, 5067-5582 and 5067-5583) and the library concentration was evaluated with the dsDNA HS assay from Qubit (Thermo Fisher, Life Technologies, Q32851). Both the library preparation and quality check were performed at the Atlantic Cancer Research Institute, Moncton, New Brunswick, Canada.

### 5.6 **Sequencing**

#### RNA-seq

Whole-genome RNA-seq and EM-seq were performed at the Next Generation Sequencing Core Facility at Atlantic Cancer Research Institute in Moncton, New Brunswick, Canada. Equimolar amounts of libraries for RNA-seq were first sequenced on the iSeq 100 instrument (Illumina) using paired-end sequencing of 2x80 to assess both library and pooling qualities. Libraries’ inputs were rebalanced following Illumina’s recommendations to ensure an equal representation of each sample. Libraries were then sequenced using NovaSeq 6000 instrument (Illumina). Samples were loaded on an SP flowcell and a paired-end sequencing of 2x80 was used. The XP workflow was utilized and samples were distributed as 12 samples on one lane for approximately 25M paired-end reads/sample.

#### EM-seq

Equimolar amounts of libraries for Enzymatic Methyl-seq were first sequenced on the iSeq 100 instrument (Illumina) using paired-end sequencing of 2x151 to assess both library and pooling qualities. Libraries’ inputs were rebalanced following Illumina’s recommendations to ensure an equal representation of each sample. Libraries were then sequenced using the NovaSeq 6000 instrument (Illumina). Samples were loaded on an S2 flowcell and a paired-end sequencing of 2x151 was used.

### 5.7 **Data analysis**

#### RNA-seq

The quality of the raw reads was verified with FastQC. Next, adapter trimming with BaseSpace™ Sequence Hub Prep platform was followed by trimming with Trim Galore (v.0.4.4, [118]) with default settings. Trimmed reads were used as input data in Salmon (v.1.4.0, [119]) to perform the pseudo alignment to the human transcriptome from Gencode (Release 37 -GRCh38.p13) and quantification. The quantification data were uploaded in the R statistical environment (v.4.0.3,[120]) using the Bioconductor package Tximeta (v.1.10, [121]). Low-expression genes were filtered, and only genes with at least more than 1 read in at least 2 samples in one of the groups were considered for further analysis. The normalization and differential expression of the count matrix [as per definition equals the number of sequence fragments that have been assigned to each gene] was performed with DESeq2 (v.1.28.1, [122]). We considered genes with p_adj_(FDR) < 0.05 and with log2FC ≥ |1| as differentially expressed (DE). The packages gplots (v.3.1.1, [123]) and ggplot2 (v.3.3.3, [124]) were used to generate both the heatmap and the volcano plot. The Reactome database (v.1.7.0, [125]) was used to identify differentially expressed pathways in humans with a t-test and p-value adjustment by the Benjamini-Hochberg formula. The ClueGO App (v.2.5.8 [126]) in Cytoscape (v.3.8.2 [127]) was used to visualize functionally grouped networks of signaling pathways relying on Reactome input. For the comparative transcriptome study, the online tool https://bioinformatics.psb.ugent.be/cgi-bin/liste/Venn/calculate_venn.htpl was used.

#### EM-seq

Fastq sequence files were generated on Illumina BaseSpace. Reads were aligned (directional) to the human (hg19 UCSC ALT-AWARE), pUC19, and lambda genome using the Dragen methylation pipeline (methyl-seq, v.3.7.5) on Illumina BaseSpace using the default Bismark parameters for methylation reporting. Alignments for spike-in (pUC19 and lambda) genomes were performed to know the conversion efficiency on the NEB EM-seq kit. Following the alignment, the data was further analyzed using the MethylKit (v.2.0.2, [128]) app on Illumina BaseSpace with a minimum CpG coverage of 5x or 10x, testing 10 to 25% methylation difference and q-value of 0.01 or 0.05.

## 6. Acknowledgements

We are grateful to Dr. Gilles Robichaud (Université de Moncton, New Brunswick, Canada) for providing cells. We thank Julie Lewis for cell transfer and Brandon Hannay for his willingness to share his expertise in HUVEC culture. NGS sequencing was performed by the Atlantic Cancer Research Institute, Moncton, New Brunswick, Canada. The authors thank Atlantic Cancer Research Institute staff members Jacynthe Lacroix, Rebecca J Shaw, Dr. Gabriel Wajnberg, Shruti Srivastava, and Simi Chacko for their help with sequencing and data analysis. We thank Dr. Gabriel Wajnberg for his critical comments on the manuscript.

## 8. Authors’ contribution

AB and VL jointly planned and designed the experiments. AB performed the experiments, the functional data analysis, and data visualization and wrote the manuscript. Both AB and VL interpreted the data, revised, and approved the manuscript.

## 10. Data availability statement

All NGS data underlying this study were deposited on database servers. RNA-seq data are accessible on NCBI’s Gene Expression Omnibus (GEO) under the super-series accession number GSE194294. EM-seq data are accessible as Sequence Read Archive (SRA) data under accession number PRJNA800079.

## Supporting information

**S1 File. RNA-seq statistics.**

**S2 File. Unfiltered processed RNA-seq data for HUVEC.** DeSeq2 was used to normalize the RNA-seq data and to analyze gene expression by comparing exposure conditions (Ctr vs. Borr). AB25.1,26.1, and 27.1 are replicates for HUVEC exposed to *Borrelia*. AB 31.1,32.1, and 33.1 are replicates for HUVEC exposed control. The sample names with suffix “.1” indicate normalized data.

**S3 File. Unfiltered processed RNA-seq data for HEK-293.** DeSeq2 was used to normalize the RNA-seq data and to analyze gene expression by comparing exposure conditions (Ctr vs. Borr). AB 28.1, 29.1, and 30.1 are replicates for HEK-293 exposed to *Borrelia.* AB 34.1,35.1, and 36.1 are replicates for HEK-293 exposed control. The sample names with suffix “.1” indicate normalized data.

**S4 File. EM-seq statistics.**

**S1 Fig. Relative log expression (RLE) plots.** Boxplots show RLE for transcriptome data obtained from human cell models HUVECs and HEK-293 cells unexposed (Ctr) or exposed (Borr) to *B. burgdorferi* strain B31 for 72 h. Analyses were performed for individual cell types (A, B) or across cells (C). Experiments were performed in triplicate as indicated by the sample names. RLE plots were generated with R package EDASeq and were used to verify data normalization with DeSeq2. Boxplots demonstrated a small deviation from zero, which is representative for the ideal distribution for normalized counts for all comparisons in this study. **S2 Fig. Heat map of differential expressed (DE) genes in HUVECs.** The cell model was unexposed (Ctr) or exposed to *B. burgdorferi* strain B31 (Borr) for 72 h in triplicates (as indicated by sample names). DeSeq2 was used to normalize the RNA-seq data and to analyze gene expression by comparing exposure conditions (Ctr vs. Borr). The list of genes was filtered for p_adj_ < 0.05 and log2FC ≥ |1|, reducing the list of samples to 69 gene loci. Shown is log2 scaled, mean-centered expression per replicate and condition in the significant DE genes of HUVECs. The column-wise dendrogram supports clustering according to the exposure (red=gBorr, black=Ctrl). Decoding of Ensembl gene IDs can be found in supplement file 3, which also lists all gene loci entered into the analysis.

**S3 Fig. Heat map of differential expressed (DE) genes in HEK-293 cells.** The cell model was unexposed (Ctr) or exposed to *B. burgdorferi* strain B31 (Borr) for 72 h in triplicates (as indicated by sample names). DeSeq2 was used to normalize the data and to analyze gene expression by comparing exposure conditions (Ctr vs. Borr). The list of genes was filtered for p_adj_ < 0.05 and log2FC ≥ |1|, reducing the list of samples to 8 significant DE genes. Shown is log2 scaled, mean-centered expression per replicate and condition in the significant DE genes of HEK-293 cells. The column-wise dendrogram supports clustering according to the exposure (red=Borr, black=Ctrl). Decoding of Ensembl gene IDs can be found in supplement file 4, which also lists all gene loci entered into the analysis.

**S4 Fig. Correlation of methylation across samples with different analytical settings.** Correlation plots for HUVECs (top line) and HEK-293 cells (bottom line) were generated with MethylKit on Illumina Basespace. Histograms displaying percent methylation per cytosine base for unexposed and *B. burgdorferi*-exposed replicates are shown diagonally per square. Numbers in the top right panels indicate Pearson correlation values of the pairwise comparison of the samples, all of which are above 0.7, indicating a strong correlation. The panels in the lower left are scatter plots of percent methylation for each sample pairing. The settings for sample comparison of *B. burgdorferi* unexposed and exposed cells varied between 10 to 25% methylation difference, minimum CpG coverage of 5 or 10x, and q-value cut-off of 0.01 or 0.05 for both HUVECs and HEK-293 cells.

